# Is there a genetic correlation between movement and immobility in field populations of a beetle?

**DOI:** 10.1101/2020.08.10.245431

**Authors:** Kentarou Matsumura, Takahisa Miyatake

## Abstract

Genetic correlations among behavioural traits are often controlled by pleiotropic genes. Many studies suggest the existence of genetic correlations among behavioural traits based on artificial selection experiments in the laboratory. However, few studies have examined whether behavioural correlations in the laboratory are maintained in the field, where natural selection works. Artificial selection experiments showed a behavioural correlation among death feigning, walking movement, and locomotor activity in the red flour beetle (*Tribolium castaneum*). This study investigated whether this behavioural correlation is observed in wild *T. castaneum* populations. We also collected beetles from various regions in Japan and investigated the geographic variation in these traits. There was geographic variation in the three behavioural traits. However, these behavioural traits were not correlated. The results suggest that the genetic correlations among behavioural traits are not maintained in the field. Therefore, the results derived from laboratory experiments may be overestimated. The same correlation between traits was not believed to arise in the field, as the indoor results may have been caused by unrealistic selection pressures. Further laboratory and field investigations are both needed.

## Introduction

Animal behaviours are often correlated with other behavioural traits due to environmental and genetic factors (Lande, 1979; Lande & Arnold, 1983; Bell, 2005). Correlations among behavioural traits by genetic factors (*i.e*., genetic correlation) are often controlled by pleiotropic genes (Lande, 1979; Lande & Arnold, 1983; Bell, 2005). If a correlation among behavioural traits has a genetic basis when natural selection favors a behaviour, other behavioural traits genetically correlated with the behavioural trait may also evolve, even if the correlated behaviours decrease fitness. This may decrease the variation in behavioural traits. That is, a genetic correlation among behavioural traits may maintain the variation in behavioural traits within a population. Therefore, studies of genetic correlations among behavioural traits are important in behavioural ecology.

Artificial selection is an experimental method that examines genetic correlations among behavioural traits (Hill & Caballero, 1992; Garland Jr & Carter, 1994). When behavioural traits respond to artificial selection for another behavioural trait, the relationship among these behavioural traits may be genetic (Falconer & Mackay, 1996). Many artificial selection experiments have examined animal behaviours. For example, risk-taking behaviour in the great tit, *Parus major* was correlated with artificial selection for exploration behaviour (van Oers, Drent, de Goede, & van Noordwijk, 2004). In the adzuki bean beetle, *Callosobruchus chinensis*, when the duration of death-feigning behaviour, which may be adaptive anti-predator behaviour, is selected artificially, flight activity responds negatively as a correlated trait (Ohno & Miyatake, 2007). However, because the pressure due to artificial selection may not occur in the field, these studies may have overestimated the correlations among behavioural traits. Therefore, comparative investigations using wild populations, not only artificial selection, are important to explain the evolution of correlations among behavioural traits.

Wild populations may be under many selection pressures compared with populations selected artificially (Mousseau, Sinervo, & Endler, 2000). If the genetic correlation among behavioural traits is controlled by pleiotropic genes, this behavioural correlation may be observed in wild populations. However, few studies have investigated behavioural correlations using both artificial selection and wild populations (but see Ohno & Miyatake, 2007).

The intensity and direction of selection pressure may differ among geographically different populations. Geographic variation in morphological and life-history traits has been observed in many species (*e*.*g*., Bergmann, 1848; Blanckenhorn & Demont, 2004; Blanckenhorn, Stillwell, Young, Fox, & Ashton, 2006). Furthermore, many studies have reported geographic variation in behavioural traits (*e*.*g*., Lankinen, 1986; Foster & Endler, 1999; Mathias, Jacky, Bradshaw, & Holzapfel, 2005; Lankinen & Forsman, 2006). Therefore, if the relationship among behavioural traits is genetic, geographic variation may also be observed in the behavioural correlation. However, few studies have investigated behavioural correlations in various wild populations.

In this study, we examined the red flour beetle *Tribolium castaneum*, a common cereal storage pest worldwide (Sokoloff, 1977). Previous studies induced artificial selection on two behavioural traits in *T. castaneum*: death-feigning behaviour and moving ability (Miyatake, Katayama, Takeda, Nakashima, Sugita, & Mizumoto, 2004; Matsumura & Miyatake, 2015). Miyatake, Katayama, Takeda, Nakashima, Sugita, & Mizumoto (2004) artificially selected death-feigning duration and established genetic strains with longer (LD) and shorter (SD) durations of death-feigning; in a predation experiment, LD individuals had a higher survival rate than the SD strain in encounters with the jumping spider, *Hasarius adansoni*. Moreover, individuals from the LD strain were significantly less mobile than the SD strain (Miyatake, Tabuchi, Sasaki, Okada, Katayama, & Moriya, 2008b). Matsumura and Miyatake (2015) artificially selected moving ability and established higher (HM) and lower (LM) mobility strains. Individuals from the HM strain had significantly shorter death feigning and greater locomotor activity than the LM strain (Matsumura, Sasaki, & Miyatake, 2016). These studies suggested that the relationship among death feigning, moving ability, and locomotor activity is genetic in *T. castaneum*. The relationship among these behavioural traits was also demonstrated in *Tribolium confusum* (Nakayama, Nishi, & Miyatake, 2010; Nakayama, Sasaki, Matsumura, Lewis, & Miyatake, 2012) and *C. chinensis* (Ohno & Miyatake, 2007; Nakayama & Miyatake, 2010) in artificial selection experiments. If these three behavioural traits are correlated genetically, such correlations may be observed in wild insect populations and geographic variation might be found in nature. Therefore, in this study, we measured the death feigning, moving ability, and locomotor activity of *T. castaneum* in wild populations at 36 locations in Japan. To investigate the role of genetic factors in the behavioural correlations, we used wild populations maintained for at least two generations after collection from the field. We investigated whether the behavioural correlation was observed in wild populations.

## Materials & Methods

### Insect

*Tribolium castaneum* was collected at 36 locations in Japan (Fig. 1). Table S1 shows the latitude and longitude of each. The northernmost is Aomori (40°89’N, 140°46’E) and the southernmost is Okinawa (26°25’N, 127°69’E) (Appendix Table A1). Collection was done in 2016 and 2017. Each beetle was reared in an incubator (Sanyo, Tokyo, Japan) maintained 25°C and 16L:8D light cycle (light on at 07:00, off at 23:00). Food is a mixture of whole meal (Nisshin Seifun, Tokyo, Japan) with brewer’s yeast (Asahi Beer, Tokyo).

**Figure 1.**
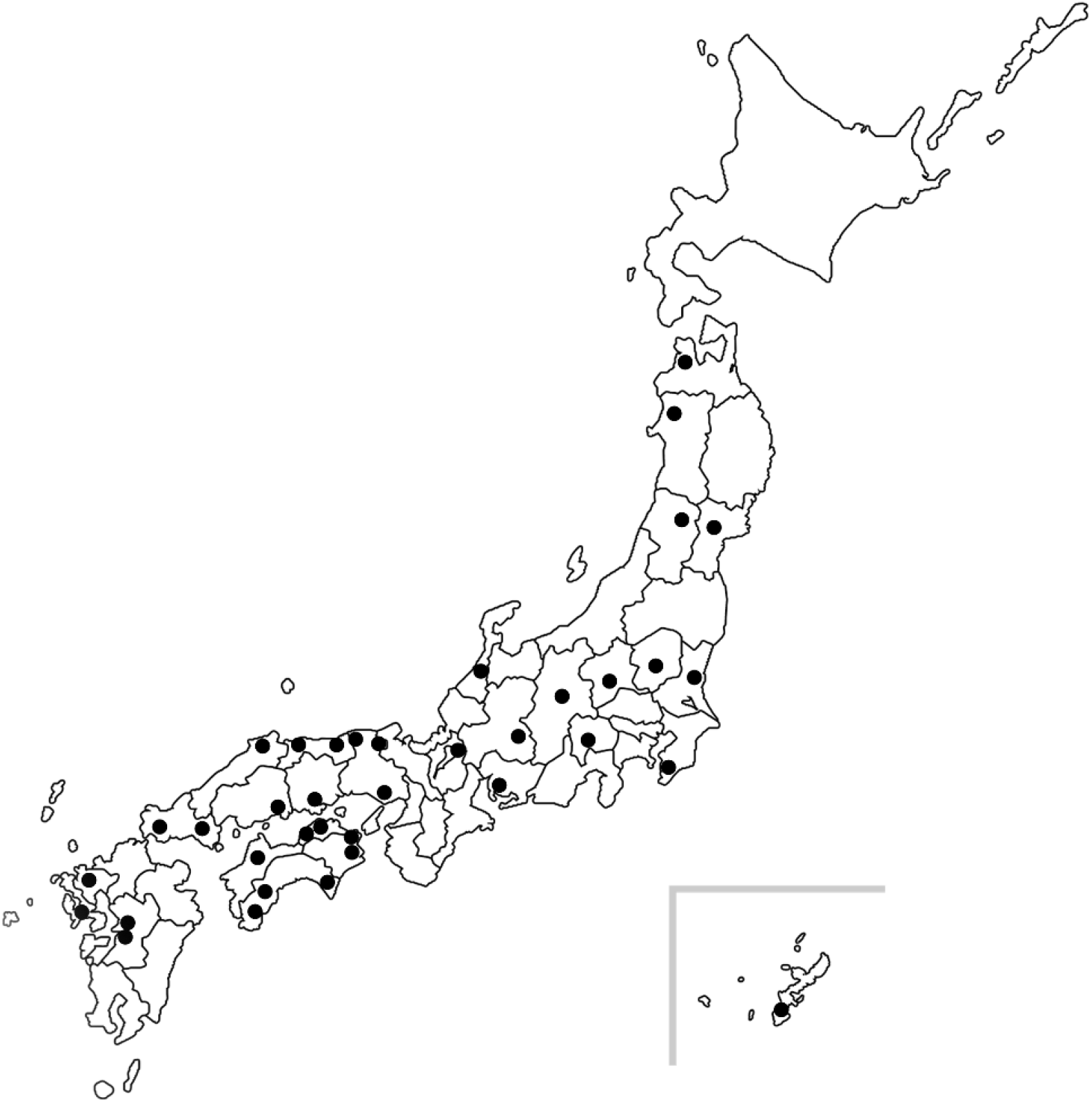
Places that captured of wild population of *T. castaneum* in Japan.

### Measurements of each behavioural trait

We measured frequency and duration of death feigning of *T. castaneum*, in accordance with Miyatake, Katayama, Takeda, Nakashima, Sugita, & Mizumoto (2004). Virgin males and females (21–28 days old) were randomly collected from each wild population. When the beetle shows death-feigning behaviour by touching the beetle’s abdomen with a stick, we measured the duration with a stopwatch (the duration was defined as the time until detecting its first visible movement). If the beetle did not show death feigning, the stimuli was repeated up to three times. Details of the methods for observation of death feigning are described in Miyatake, Katayama, Takeda, Nakashima, Sugita, & Mizumoto (2004).

Walking distance as the moving ability of *T. castaneum* was measured using an image tracker system (Digimo, Osaka, Japan), in accordance with Matsumura and Miyatake (2015). Virgin beetles (21–28 days old) were collected from each population and measured walking distance. The moving behaviour were recorded for 30 min, and measure the walking distance of each beetle on the recorded image, we used analysis software (2D-PTV Ver. 9.0, Digimo, Osaka, Japan). Details of the methods for measurement of moving ability are described in Matsumura and Miyatake (2015).

To measurement of locomotor activity, virgin males and females (21–28 days old) were randomly collected from each wild population, and measured locomotor activity used by an infrared actograph system. When the beetle passed the midpoint of the dish, the infrared light between emitter and detector device (E3R-5E4/R2E4DS30E4; Omron, Kyoto, Japan) was interrupted. We measured the number of interruptions of the infrared light for 24 h as locomotor activity of the beetle. To remove artificial effect on behavior, we removed first 2 h data (i.e., we used 22 h data for statistical analysis). Details of the methods for measurement of locomotor activity are described in Matsumura, Sasaki, & Miyatake (2016).

### Statistical analysis

Death-feigning duration (+ 1 s) and moving ability (+ 1 mm) were analyzed using a generalized linear model (GLM) with gamma distribution, and population, sex, and the interaction between population and sex as experimental variables. The frequencies of death feigning and locomotor activity were analyzed by GLMs with binomial and Poisson distributions, respectively. To analyze the relationship between duration of death feigning and moving ability within individual level, we used GLM with gamma distribution, and duration of death feigning as a dependent variable, and moving ability and population as experimental variables. Because individuals used measurement of locomotor activity are differed with individuals used measurement of death feigning and moving ability, we did not analysis of relationship between death feigning and locomotor activity, and moving and locomotor activity within individual. To analyze the relationship between death feigning, moving ability, and locomotor activity inter populations, we used Spearman’s rank correlation coefficient for mean values of three behavioural traits in each wild population. To analyze the effects of latitude and longitude on mean values of each behavioural trait, we used analysis of covariance (ANCOVA) with population, sex, and the interaction between the two as covariates. All analyses were done using R ver. 3.4.3 (R Core Team, 2017).

### Ethical Note

The laboratory population of *T. castaneum* used in this study have maintained at Okayama University for over 15 years. This population has been maintained on whole meal flour with yeast (see Miyatake et al. 2004). We reared this population at 25 °C, which resemble natural conditions for this insect. All animals in the study were handled more carefully. The use of these animals conforms to the Animal Ethics Policy of Okayama University.

## Results

Figure 2 shows the duration and frequency of death-feigning behaviour. Figure 3 shows the moving ability and locomotor activity. Table 1 shows the mean values of each behaviour. Death-feigning duration differed significantly among wild populations, but not the sexes (Fig. 2a, Table 2). The frequency of death feigning also differed significantly among wild populations, and was significantly more frequent in males (Fig. 2b, Table 2). Moving ability differed significantly among wild populations, but not the sexes (Fig. 3a, Table 2). Locomotor activity differed significantly among wild populations, and females were significantly more active than males (Fig. 3b, Table 2). The interaction between wild population and sex was not significant in any behavioural trait (Table 2). Appendix Figure A1 shows the relationship between latitude and each behavioural trait. There were no significant associations between latitude and each behavioural trait (Table 3). Appendix Figure A2 shows the relationship between longitude and each behavioural trait. There were no significant associations between longitude and each behavioural trait (Table 3). Duration of death feigning did not show significantly correlated with moving ability within individual level (*χ*^2^_1,1044_ = 3.80, *p* = 0.05112). In inter population level, no behaviour was significantly correlated with another (Table 4).

**Table 1.** Mean and SE of each trait.

| Trait | Male |  | Female |  |
| --- | --- | --- | --- | --- |
| | Mean ± SE | *N* | Mean ± SE | *N* |
| Duration of death feigning (s) | 116.35 ± 5.26 | 1064 | 120.81 ± 6.31 | 1022 |
| Frequency of death feigning | 0.94 ± 0.01 | 1064 | 0.92 ± 0.01 | 1022 |
| Walking distance (mm) | 214.11 ± 17.49 | 533 | 184.63 ± 14.52 | 533 |
| Locomotor activity | 95.52 ± 7.09 | 614 | 216.62 ± 19.82 | 605 |

**Table 2.**
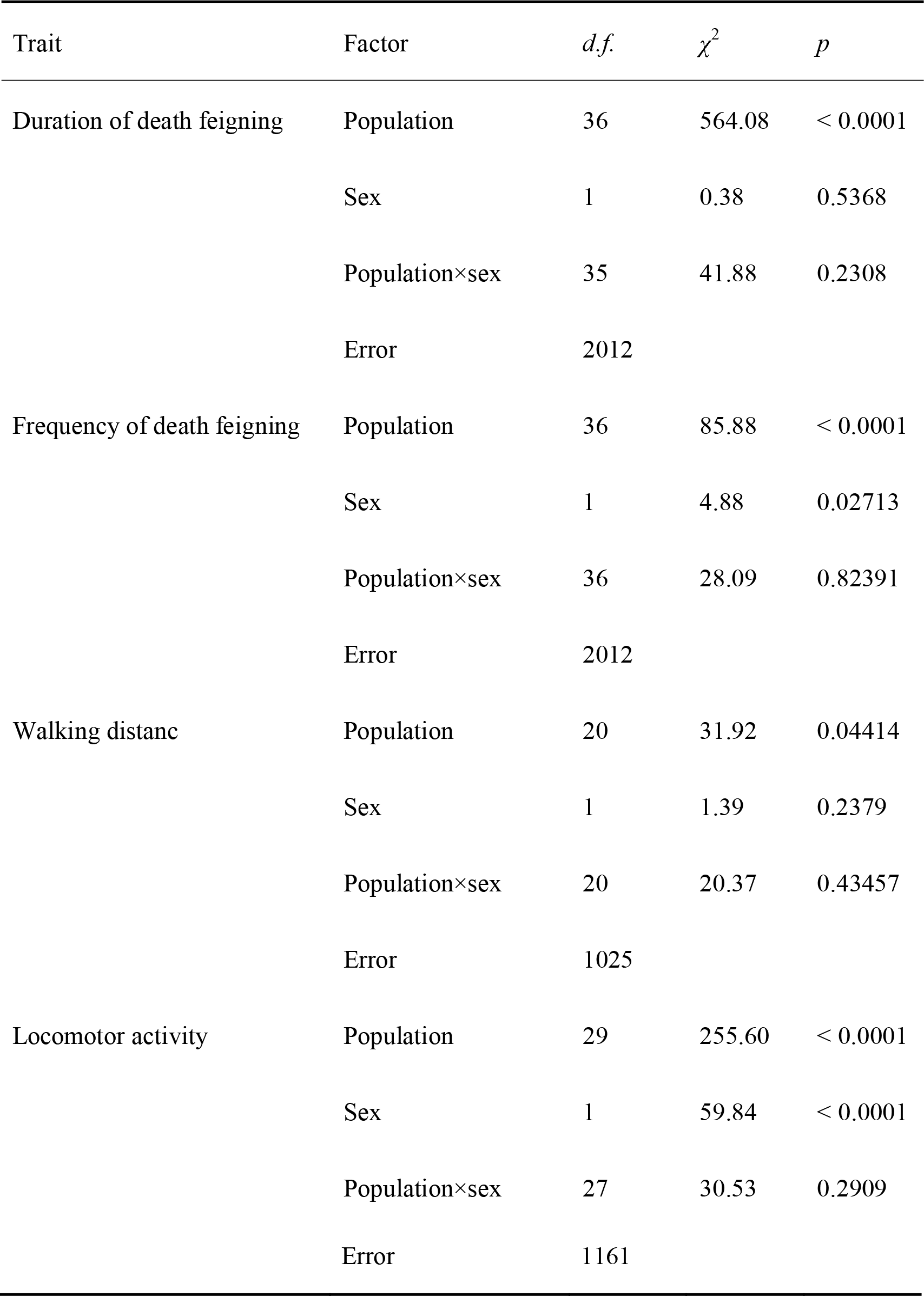
Results of GLM for effects of population and sex on death feigning, walking distance, and locomotor activity.

**Table 3.** Results of mixed ANOVA for effects of latitude, longitude and sex on each behavioural traits.

| Trait | Factor | *d.f.* | *F* | *P* |
| --- | --- | --- | --- | --- |
| Death feigning | Latitude | 1 | 0.05 | 0.8199 |
|  | Longitude | 1 | 1.87 | 0.1803 |
|  | Sex | 1 | 0.26 | 0.6128 |
|  | Error | 2082 |  |  |
| Mobility | Latitude | 1 | 0.01 | 0.9276 |
|  | Longitude | 1 | 0.00 | 0.9674 |
|  | Sex | 1 | 1.74 | 0.1880 |
|  | Error | 1062 |  |  |
| Activity | Latitude | 1 | 0.01 | 0.9143 |
|  | Longitude | 1 | 0.15 | 0.6993 |
|  | Sex | 1 | 40.80 | < 0.0001 |
|  | Error | 1217 |  |  |

**Table 4.**
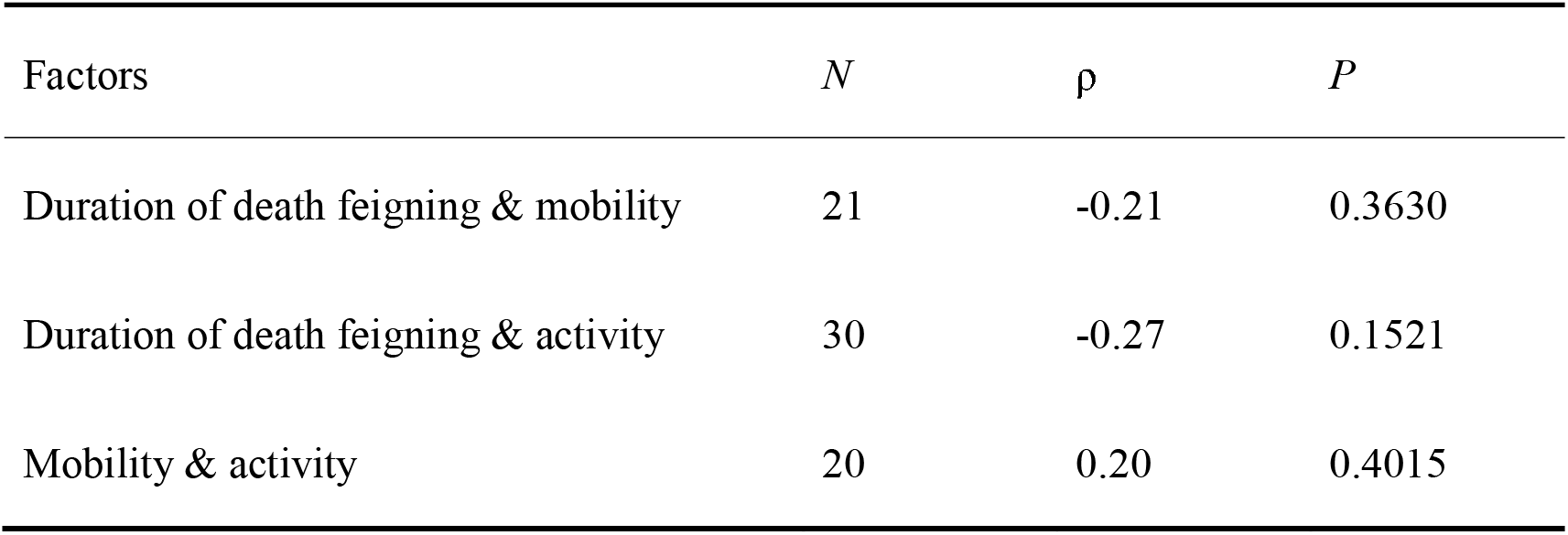
Results of Spearman’s rank correlation coefficient for mean values in each behavioural trait.

**Figure 2.**
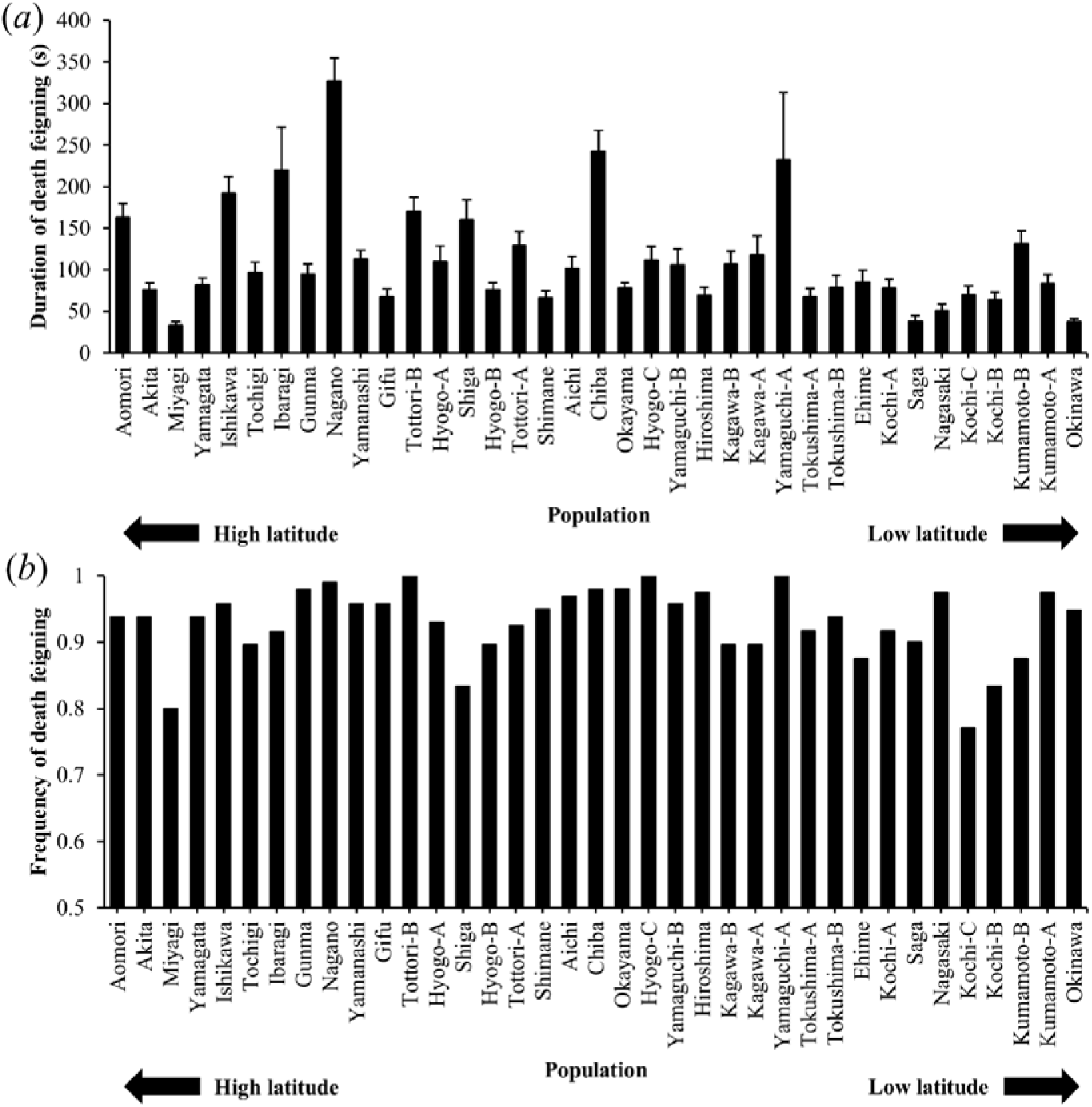
Duration (a) and frequency (b) of death-feigning behaviour in each population. Error bars show SE.

**Figure 3.**
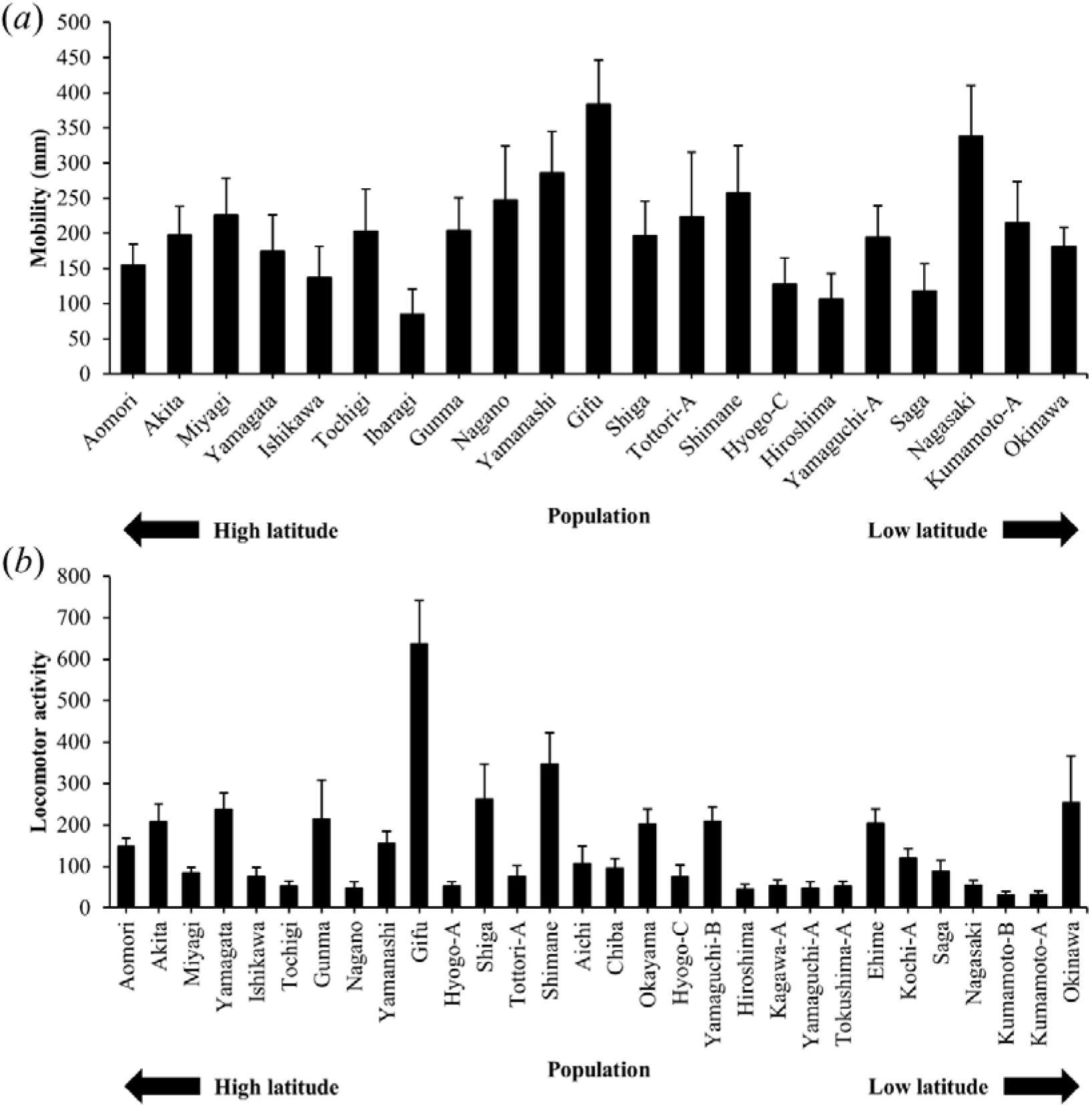
Mobility (a) and locomotor activity (b) in each population. Error bars show SE.

## Discussion

This study found geographic variation in death-feigning, moving ability, and locomotor activity among 36 wild populations collected in Japan (Figs. 2, 3). Because we used each population maintained at the laboratory for at least two generations, these behavioural differences among populations may be caused by genetic factors, rather than environmental or maternal factors. Moreover, the geographic variation suggests that the selection pressures on the three behaviours differ among wild populations. However, although previous studies suggested genetic correlations among the three behaviours in artificial selection experiments, we did not find any correlations among the three behaviours in the wild populations (Fig. 4). This suggests that the intensity of the relationships among these behavioural traits is lower in the field, even if the genetic correlations were shown by artificial selection.

**Figure 4.**
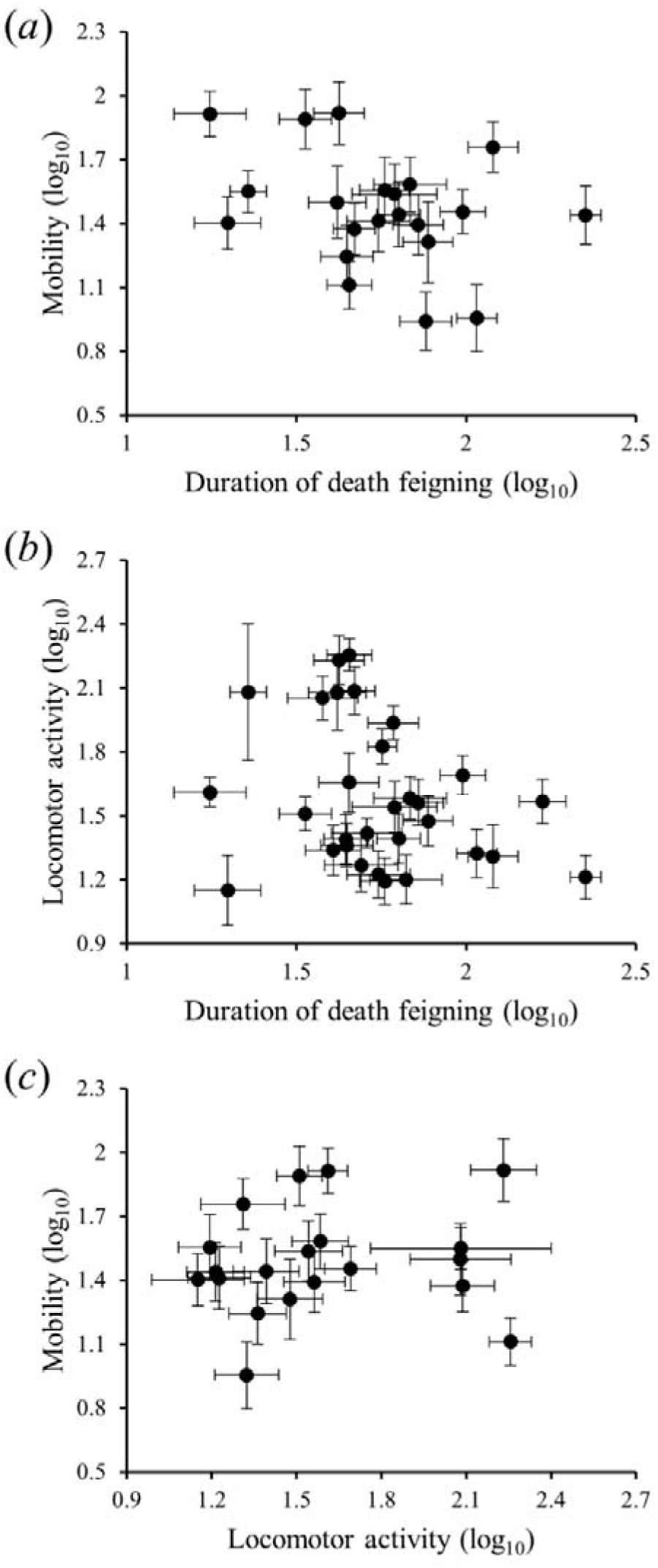
Relationship between duration of death feigning and walking speed or locomotor activity. Each point shows mean value of each population. Error bars show SE.

First, we discuss the results for the death-feigning behaviour. Prohammer and Wade (1981) reported geographic variation in death-feigning behaviour among populations from Spain, Japan, and USA. Our results are in accord regarding geographic variation in the duration of death feigning (Prohammer & Wade, 1981). We also found that geographic variation in death feigning was seen over a relatively narrow range compared with Prohammer & Wade (1981).

We considered three hypotheses to explain the death-feigning behaviour results. The first is the difference in predation pressure on each wild population. Death feigning is an adaptive anti-predator behaviour (Humphreys & Ruxton, 2018). However, the optimum death-feigning duration may be affected by predator type or density. In the great tit *Parus major*, the egg ejection rate showed geographic variation, and this pattern matched the geographic pattern of parasitism risk (Liang et al., 2016). In *T. castaneum*, individuals with longer death-feigning duration increase their survival rate in encounters with predators (Miyatake, Katayama, Takeda, Nakashima, Sugita, & Mizumoto, 2004). Similarly, the optimum death-feigning duration may be altered by the density of predators. For example, individuals with longer death-feigning duration may increase the survival rate in places under higher predation pressure, whereas individuals with a shorter duration of this behaviour may increase foraging or reproductive success at places with lower predation pressure (Nakayama & Miyatake, 2010a, b). Moreover, differences in the type of predation may affect the evolution of this behaviour (Honma, Oku, & Nishida, 2006). For example, individuals with longer death-feigning duration may increase their survival rate when encountering active-hunting predators, whereas they may decrease the survival rate when encountering sit-and-wait predators (Honma, Oku, & Nishida, 2006). Therefore, in locations with sit-and-wait predators, shorter death-feigning duration may evolve. Populations with longer death-feigning duration might suffer predation pressure by active hunting predators, whereas populations with shorter death feigning might suffer predation pressure by sit-and-wait predators. Ohno & Miyatake (2007) also reported geographic variation in the duration of death feigning in *C. chinensis*, and suggested that this geographic variation was a result of differences in predation pressure among these wild populations. Future studies should investigate the density and type of predators.

The second hypothesis is the effects of prey density. In *T. castaneum*, the optimal death-feigning duration may depend on the conspecific density. Miyatake, Nakayama, Nishi, & Nakajima (2009) reported that beetles with longer death-feigning duration had a higher survival rate in the presence of non-feigners or prey of a different species, compared to when alone, confirming the selfish-prey hypothesis. The prey density might have important effects on death-feigning behaviour. Conspecific or heterospecific density in populations with longer death-feigning durations may be higher, whereas it may be lower in populations with shorter death-feigning durations. Studies need to investigate the effects of population density.

The third hypothesis is founder effects on the behaviour. The founder effect, *i*.*e*., the loss of genetic variation when a new population is established by a small number of individuals (Templeton, 1980), is found in animal behaviour (Suarez, Tsutui, Holway, & Case, 1999). In populations with longer death-feigning duration, populations may have been established by individuals with longer death-feigning durations. This hypothesis is based on the supposition that gene flow is infrequent among *T. castaneum* populations. Although a study reported a bottleneck in a wild population (Semeao, Campbell, Beeman, Lorenzen, Whitworth, & Sloderbeck, 2012), another suggested that gene flow often occurs over a wide range in *T. castaneum* (Ridley, Hereward, Daglish, Raghu, Collins, & Walter, 2011). Additional studies should investigate genetic differences among wild populations.

Moving ability and locomotor activity also showed geographic variation among wild populations in *T. castaneum*. A previous study revealed that *T. castaneum* with genetically higher moving ability had a lower survival rate when predators were present (Matsumura & Miyatake, 2015). Similarly, individuals with higher locomotor activity are considered at increased risk of predation in many animals (Sih, Bell, & Johnson, 2004; Réale, Reader, Sol, McDougall, & Dingemanse, 2007). Therefore, predation pressure on moving ability and locomotor activity may also differ among wild populations, as with death-feigning behaviour. Moreover, moving ability affects reproductive success, and there was a trade-off between survival rate and reproductive success between strains selected for higher and lower moving ability in *T. castaneum* (Matsumura & Miyatake, 2015, 2018a; Matsumura, Archer, Hosken, & Miyatake, 2019). Therefore, these behavioural traits may be affected by balancing selection between predation avoidance and reproduction among wild populations. Moving ability seems to be similar to locomotor activity and vice versa. Nevertheless, these behaviours were not significantly correlated. These results suggest that these behaviours evolved independently in the field. In the parasitoid wasp *Leptopilina heterotoma*, geographic variation was found in locomotor activity (Fleury, Allemand, Fouillet, & Boulétreau, 1995). Furthermore, some insect studies reported that the circadian rhythm of locomotor activity showed geographic variation, with a clear rhythm at lower latitudes and no rhythmic activity at higher latitudes (*e*.*g*., Fleury, Allemand, Fouillet, & Boulétreau, 1995; Joshi, 1999). Additional studies should investigate the circadian rhythm of *T. castaneum* in each wild population.

Although this study revealed geographic variation in three behavioural traits, these behavioural traits did not show latitudinal or longitudinal clines (Appendix Figs. A1, A2). In the medaka *Oryzias latipes* complex, courtship behaviour by males and female preference for males showed latitudinal variation such that populations from lower latitudes showed greater intensity of these behaviours (Fujimoto, Miyake, & Yamahira, 2015). These results suggest that because the reproductive season is relatively shorter at lower latitudes, a latitudinal cline develops in the operational sex ratio among wild populations, which ultimately results in a latitudinal cline in sexual selection pressures (Fujimoto, Miyake, & Yamahira, 2015). Therefore, the intensity or direction of selection pressure on the behavioural traits may differ among wild populations from various latitudes. Moreover, some studies reported that individuals had a significantly longer death-feigning duration under lower temperatures than under higher temperatures in other insect species (Holmes, 1906; Miyatake, Okada, & Harano, 2008a). However, we did not find a latitudinal or longitudinal cline in death-feigning behaviour. That is, the effects of temperature on death feigning may be relatively smaller in *T. castaneum*, at least, within the temperature range examined in this study. Furthermore, moving ability and locomotor activity did not show latitudinal or longitudinal clines, which also suggested that environmental factors, such as temperature, do not affect these behavioural traits in this beetle.

Although the three behavioural traits showed geographic variation among the wild populations, we did not find a significant correlation among these behaviours (Fig. 4). Some artificial selection studies in laboratories suggested a genetic correlation among death-feigning, moving ability, and locomotor activity in some insects (Ohno & Miyatake, 2007; Miyatake, Tabuchi, Sasaki, Okada, Katayama, & Moriya, 2008b; Nakayama & Miyatake, 2010; Nakayama, Nishi, & Miyatake, 2010; Nakayama, Sasaki, Matsumura, Lewis, & Miyatake, 2012; Matsumura, Fuchikawa, & Miyatake, 2017). Therefore, we hypothesized that there would be a correlation among these behavioural traits in wild populations. However, we did not find a significant correlation among these behaviours. The results suggest that the intensity of the genetic correlations among each behaviour may be relatively lower in *T. castaneum*. Previous studies reported that moving ability and locomotor activity showed correlated responses to artificial selection for death-feigning behaviour for over 10 generations (Ohno & Miyatake, 2007; Miyatake, Tabuchi, Sasaki, Okada, Katayama, & Moriya, 2008b; Nakayama, Nishi, & Miyatake, 2010). Moreover, death-feigning and locomotor activity showed correlated responses to artificial selection for moving ability for more than 15 generations (Matsumura, Fuchikawa, & Miyatake, 2016). If the intensity of the genetic correlations among these behaviours is relatively low, these correlations may be difficult to observe in the field. Alternatively, the artificial selection pressure may be of abnormal intensity compared with the field. For example, although beetles from strains selected artificially for longer death-feigning duration feigned death for up to 1,000 seconds (Miyatake, Katayama, Takeda, Nakashima, Sugita, Mizumoto, & Miyatake, 2004; Matsumura & Miyatake, 2018b), few beetles feigned death this long in the wild populations (Fig. 2a). Another study reported that the duration of death feigning was correlated with flight activity within and among wild populations in *C. chinensis* (Ohno & Miyatake, 2007). Therefore, the intensity of the relationship between behaviours may differ among behaviours or species. Similar investigations of other behaviours or species are required.

In conclusion, a genetic correlation seen in artificial selection experiments in the laboratory among behavioural traits may not always be observed in the field. This is because the selection pressure due to artificial selection may be abnormal. This suggests that artificial selection may lead to overestimation of the results. Therefore, to study the evolution of genetic correlations in behavioural traits, behavioural ecologists should examine the relationships among behavioural traits using both artificial selection and wild populations.

## Acknowledgement

We thanks to Mr. Yusuke Tsushima and Kohei Nakao for collection of *T. castaneum* in Aomori and Hyogo prefectures. This work was supported by a grant from the Japan Society for the Promotion of Science KAKENHI 26291091, 16K14810, 17H05976 and 18H02510 to TM.

## Appendix

**Table A1.** Name, latitude, longitude, and sample size of each behavioural traits of each place that captured of populations of *T. castaneum*.

| Population | Latitude | Longitude | Sample size |  |  |
| --- | --- | --- | --- | --- | --- |
|  |  |  | Death feigning | Walking speed | Locomotor activity |
| Aomori | 40.8909 | 140.4552 | 80 | 77 | 80 |
| Akita | 40.1825 | 140.0518 | 80 | 79 | 20 |
| Miyagi | 38.5726 | 140.8723 | 40 | 40 | 64 |
| Yamagata | 38.4264 | 140.3856 | 80 | 78 | 20 |
| Ishikawa | 36.6707 | 136.7276 | 96 | 48 | 42 |
| Tochigi | 36.5747 | 139.8776 | 48 | 48 | 42 |
| Ibaragi | 36.3746 | 140.4169 | 95 | 48 | - |
| Gunma | 36.3546 | 139.1299 | 48 | 48 | 64 |
| Nagano | 35.9904 | 137.9795 | 96 | 47 | 44 |
| Yamanashi | 35.5952 | 138.5768 | 48 | 47 | 64 |
| Gifu | 35.5583 | 137.4489 | 48 | 48 | 64 |
| Tottori-B | 35.5357 | 134.3827 | 48 | - | - |
| Hyogo-A | 35.4942 | 134.5642 | 43 | - | 64 |
| Shiga | 35.4862 | 136.2289 | 48 | 48 | 64 |
| Hyogo-B | 35.4649 | 134.9447 | 48 | - | - |
| Tottori-A | 35.4216 | 133.7830 | 40 | 40 | 24 |
| Shimane | 35.2467 | 132.5565 | 40 | 38 | 22 |
| Aichi | 35.0232 | 136.9688 | 96 | - | 42 |
| Chiba | 35.0029 | 139.9148 | 48 | - | 43 |
| Okayama | 34.7247 | 133.6436 | 100 | - | 76 |
| Hyogo-C | 34.6903 | 134.9988 | 40 | 40 | 24 |
| Yamaguchi-B | 34.5033 | 131.4720 | 48 | - | 64 |
| Hiroshima | 34.5006 | 133.3735 | 40 | 40 | 30 |
| Kagawa-B | 34.3208 | 133.9018 | 48 | - | - |
| Kagawa-A | 34.2553 | 133.7846 | 48 | - | 32 |
| Yamaguchi-A | 34.2240 | 131.8132 | 40 | 40 | 20 |
| Tokushima-A | 34.0737 | 134.3950 | 48 | - | 32 |
| Tokushima-B | 33.9734 | 134.6358 | 48 | - | - |
| Ehime | 33.9224 | 133.0625 | 48 | - | 32 |
| Kochi-A | 33.4298 | 133.9903 | 48 | - | 32 |
| Saga | 33.2845 | 130.0928 | 40 | 40 | 32 |
| Nagasaki | 33.0817 | 129.8691 | 40 | 40 | 34 |
| Kochi-C | 33.0401 | 133.0547 | 48 | - | - |
| Kochi-B | 32.9847 | 132.9482 | 48 | - | - |
| Kumamoto-B | 32.8051 | 130.8083 | 48 | - | 20 |
| Kumamoto-A | 32.7974 | 130.7815 | 40 | 39 | 24 |
| Okinawa | 26.2459 | 127.6915 | 96 | 93 | 4 |

**Figure A1.**
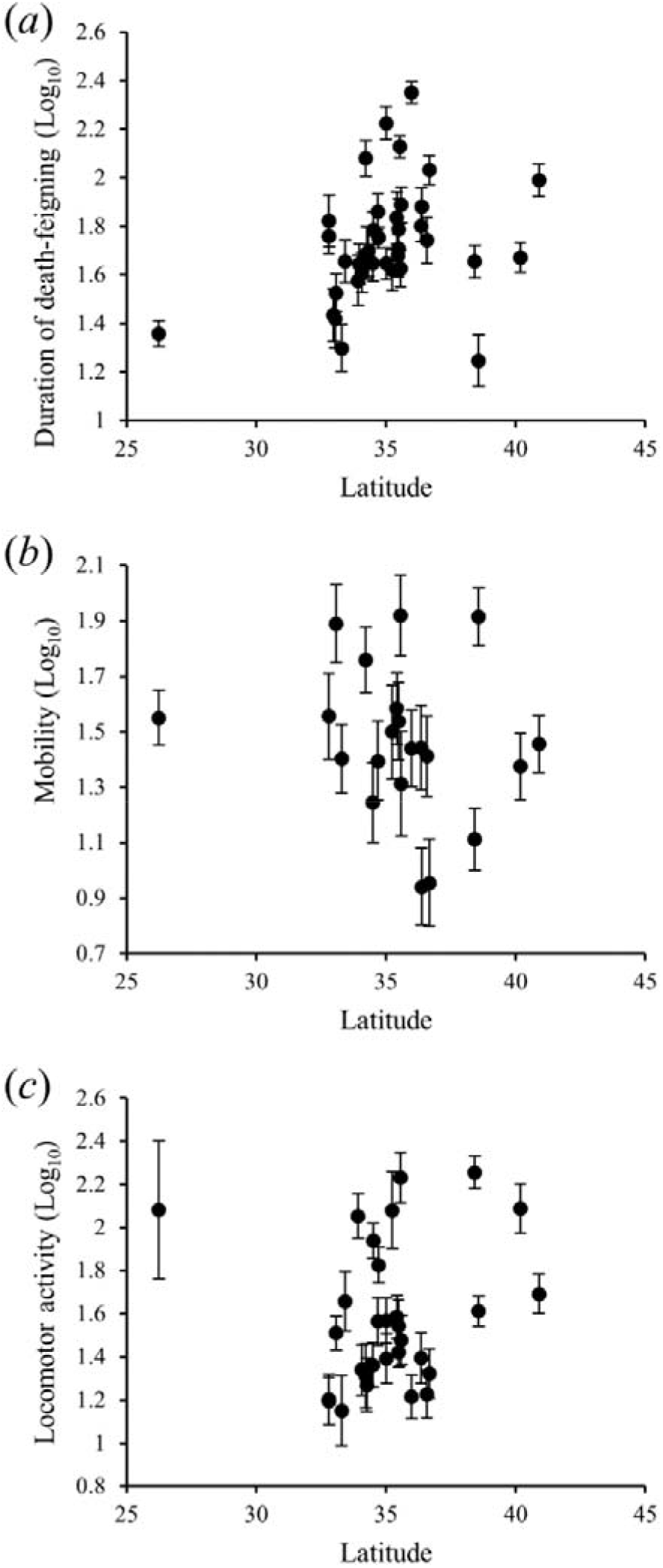
Relationship between latitude and each behavioural trait (a: death-feigning, b: mobility, c: locomotor activity). Error bars show SE.

**Figure A2.**
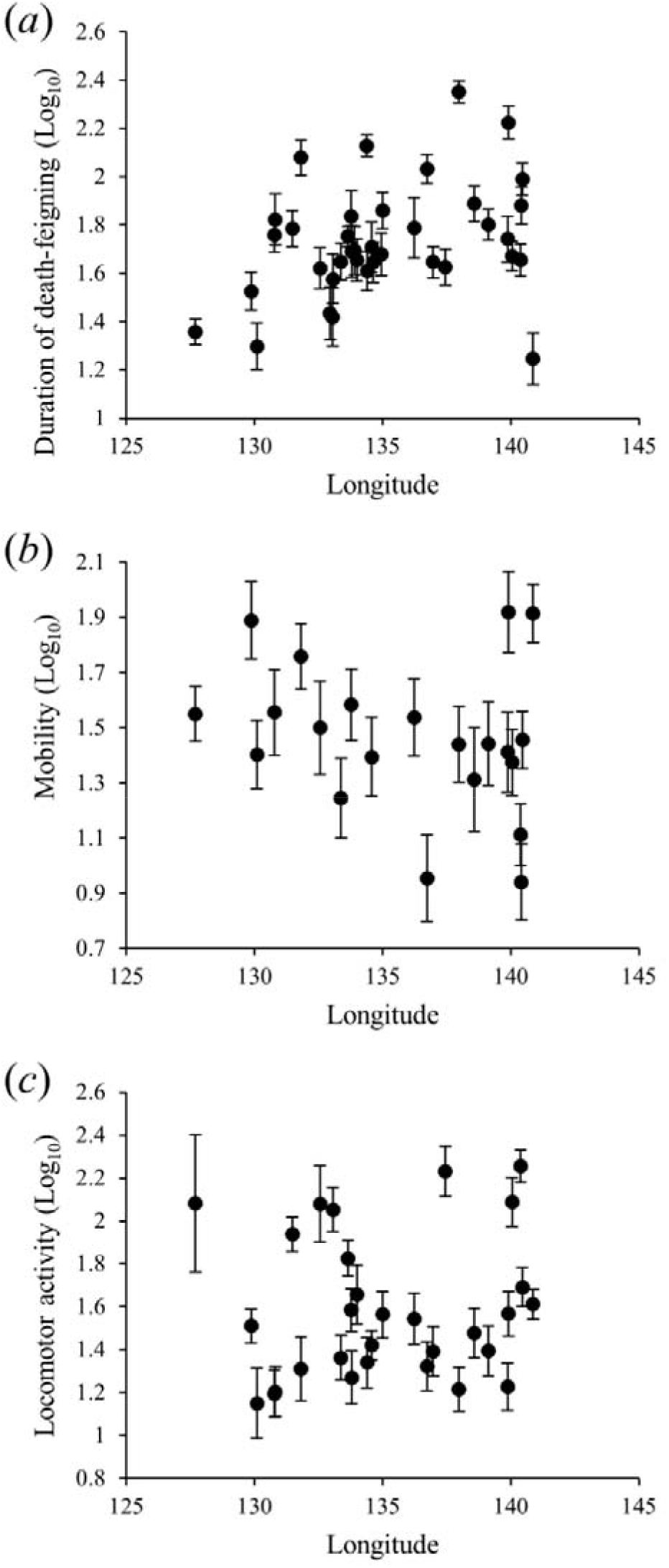
Relationship between longitude and each behavioural trait (a: death-feigning, b: mobility, c: locomotor activity). Error bars show SE.

